# PEPPI: Whole-proteome protein-protein interaction prediction through structure and sequence similarity, functional association, and machine learning

**DOI:** 10.1101/2021.12.02.470917

**Authors:** Eric W. Bell, Jacob H. Schwartz, Peter L. Freddolino, Yang Zhang

## Abstract

Proteome-wide identification of protein-protein interactions is a formidable task which has yet to be sufficiently addressed by experimental methodologies. Many computational methods have been developed to predict proteome-wide interaction networks, but few leverage both the sensitivity of structural information and the wide availability of sequence data. We present PEPPI, a pipeline which integrates structural similarity, sequence similarity, functional association data, and machine learning-based classification through a naïve Bayesian classifier model to accurately predict protein-protein interactions at a proteomic scale. Through benchmarking against a set of 798 ground truth interactions and an equal number of noninteractions, we have found that PEPPI attains 4.5% higher AUROC than the best of other state-of-the-art methods. As a proteomic-scale application, PEPPI was applied to model the interactions which occur between SARS-CoV-2 and human host cells during coronavirus infection, where 403 high-confidence interactions were identified with predictions covering 73% of a gold standard dataset from PSICQUIC and demonstrating significant complementarity with the most recent high-throughput experiments. PEPPI is available both as a webserver and in a standalone version and should be a powerful and generally applicable tool for computational screening of protein-protein interactions.

## 1. Introduction

The biological function of many proteins is conferred through their interactions with other proteins. Therefore, to fully understand the function of each protein in an organism, one must first attain a comprehensive network of the protein-protein interactions (PPIs) that occur within the cell. The discovery of critical interactions within this interaction network, or “interactome”, can lead to drug development[1] or protein engineering[2, 3] targeting these interactions. However, many of these experiments do not guarantee that the interactions detected are, in fact, direct physical contacts between the proteins; some of the earliest databases for PPI prediction involve features that assert only a functional association between proteins[4]. While these databases can be useful for prediction of physical interactions, as all physical interactions are functionally associated, the converse is not true; many biological applications, such as drug target discovery, require knowledge of which proteins come into physical contact. The methods for elucidating these direct physical interactions are at present either prohibitively costly for whole-proteome analysis (such as structure solving or crosslinking mass spectrometry) or are too susceptible to errors (such as yeast-two hybrid[5]). As an alternative, computational methods can be used to model proteome-wide interactions, as well as refine existing interaction datasets.

One of the most straightforward methods of computational interaction prediction is to determine whether the query protein pair is similar to an already known interaction. Many programs directly leverage sequence similarity for this purpose because the sequence comparison operation is quick and sequence data is plentiful[6, 7]. However, since structure is more evolutionarily conserved than sequence, structural similarity is much more effective at detecting distantly similar PPIs; methods which leverage this structural information[8–11] grow more powerful as modern structural biology methods such as cryo-EM facilitate the solving of complicated protein complex structures, and as computational approaches offer improved accuracy in predicting the folds of individual proteins[12]. In addition, structures provide a clear ground truth as to whether two proteins interact physically; if a solved structure of the interaction exists, the proteins are likely to interact *in vivo*. Therefore, an effective similarity-based program should consider both structural and sequence similarity.

Another common method for PPI prediction is the application of machine learning-based classifiers. In the earlier days of machine learning, the major novelty of machine learning-based PPI predictors was in how they extracted features from the input amino acid sequences in order to create a fixed-length vector that could be utilized in standard machine learning algorithms, such as the conjoint triad method[13] or autocorrelation 14]. As more modern deep learning techniques became available, improved PPI predictions were achieved with features solely from evolutionary profiles[15] or even based on sequence alone[16, 17]. Despite these initial studies into deep learning, to our knowledge these methods have yet to demonstrate success in a species agnostic context; more commonly, they are benchmarked by either combining a few species-specific datasets[16] or by training on one species and testing on another[15, 17].

Here, we present a Pipeline for the Extraction of Predicted Protein-protein Interactions (PEPPI), which offers high-accuracy PPI predictions through a consensus of sequence and structural similarity, functional association, and neural network classification. While the source code for PEPPI can be found at https://github.com/ewbell94/PEPPI, an online webserver implementation of this pipeline can be found at https://zhanggroup.org/PEPPI/, which allows users to create PPI predictions from sequences alone. We additionally present an application of PEPPI to make predictions of the inter-species interactome between human host cells and SARS-CoV-2. Through the following benchmarks and examples, we demonstrate that PEPPI is a useful tool for predicting both pairwise and systems-level PPIs.

## 2. Results

### 2.1 Pipeline Overview and Module Cross-Validation

PEPPI is a protein-protein interaction prediction pipeline which takes in a pair of query sequences and quantifies their likelihood of interaction as a natural log-transformed likelihood ratio (log(LR)) through a consensus of five independent prediction modules (Figure 1). This consensus is determined by a naïve Bayesian classifier model trained on a set of 800 high-confidence interactions from IntAct[18] and 800 curated non-interactions from the Negatome 2.0 database[19] (see Supplementary Methods).

**Figure 1.**
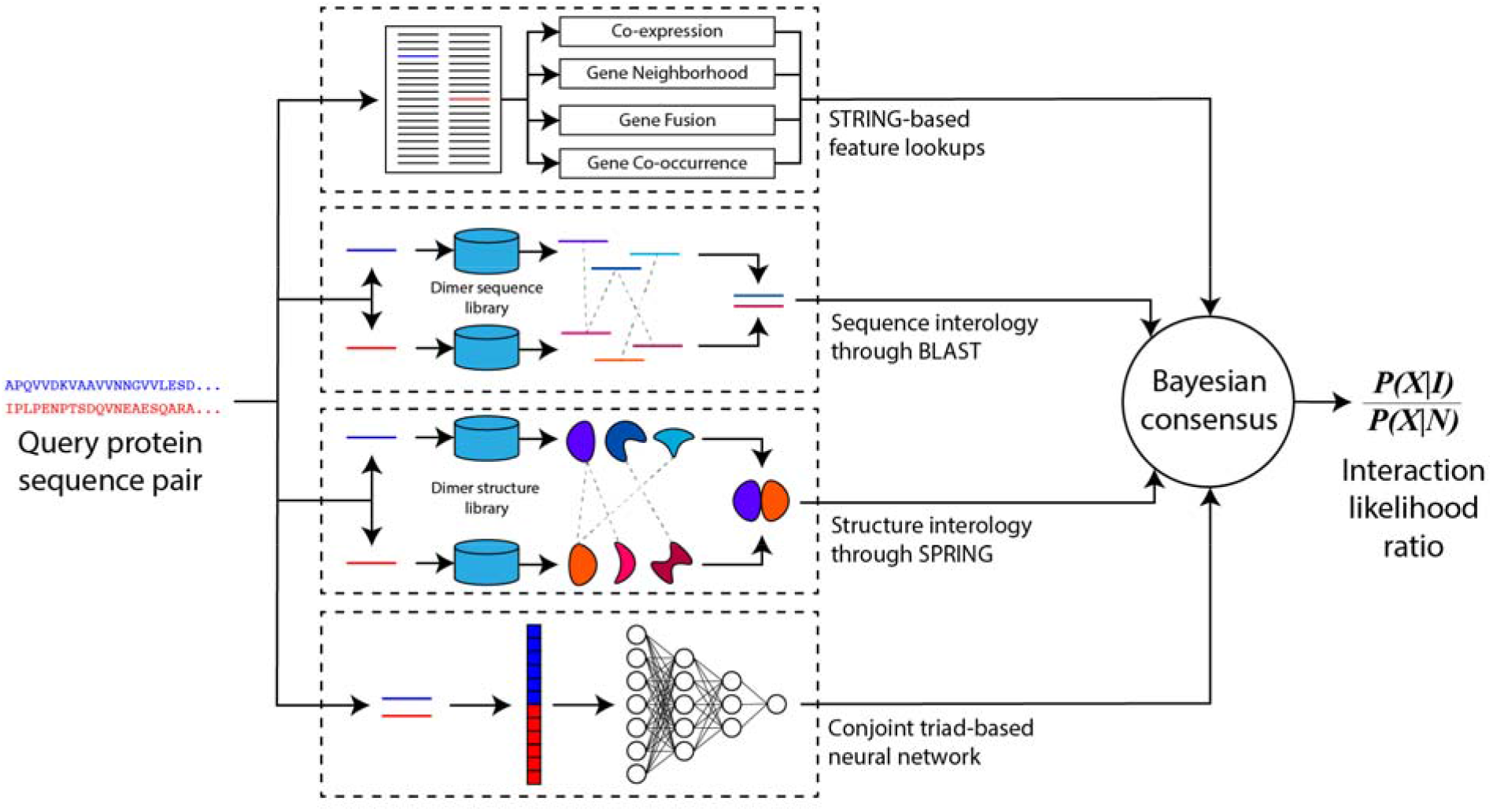
An illustration of the PEPPI pipeline. This pipeline functions by analyzing a pair of input sequences via a series of independent modules, including structure similarity, sequence similarity, neural network classification, and functional association data. These modules are combined using a naïve Bayesian consensus classifier, which provides the final interaction score as a log-likelihood ratio.

Figure 2a presents the 10-fold cross validation performance of each individual module and the full pipeline on this training set. The best-performing individual modules are SPRING and SEQ, which implement structure and sequence-based similarity approaches, respectively. The SPRING module uses the dimeric threading program SPRING[20] to identify dimer structure templates out of a database of interacting proteins extracted from the PDB, while SEQ uses BLAST[21] sequence searching to identify similar interactions in a database of direct interactions identified by high-throughput experiment (HTE) data. These homology-based modules will perform well for any cases which have homologous similarity to existing interactions, which is the case for many true interactions. The next best-performing module is the neural network-based CT module, which transforms the input amino acid sequences into a fixed-length vector according to the conjoint triad method[13] and classifies the resulting vector through a neural network model. This module helps PEPPI retrieve true positive predictions in case there is only loose homologous similarity to existing interactions. The STRING module, a module which extracts various query functional association features from the STRING database[22], performs relatively poorly on its own; this is expected because of its focus on functional association data instead of physical interaction data and because of its tendency to provide no data if the interaction is not located in STRING. SPRINGNEG has a nearly identical methodology to SPRING but it performs least well because it searches through a database of non-interacting protein structures and thus a hit in this database will only lower the interaction score because by design it only provides information to filter out functionally associated noninteractions. STRING also gives insight into non-interactions, as the training data revealed that a high co-expression value is more likely to belong to a false positive functional association than a direct physical interaction. Overall, the combination of all modules clearly outperforms any single module, demonstrating that the modules are complimentary in classification.

**Figure 2.**
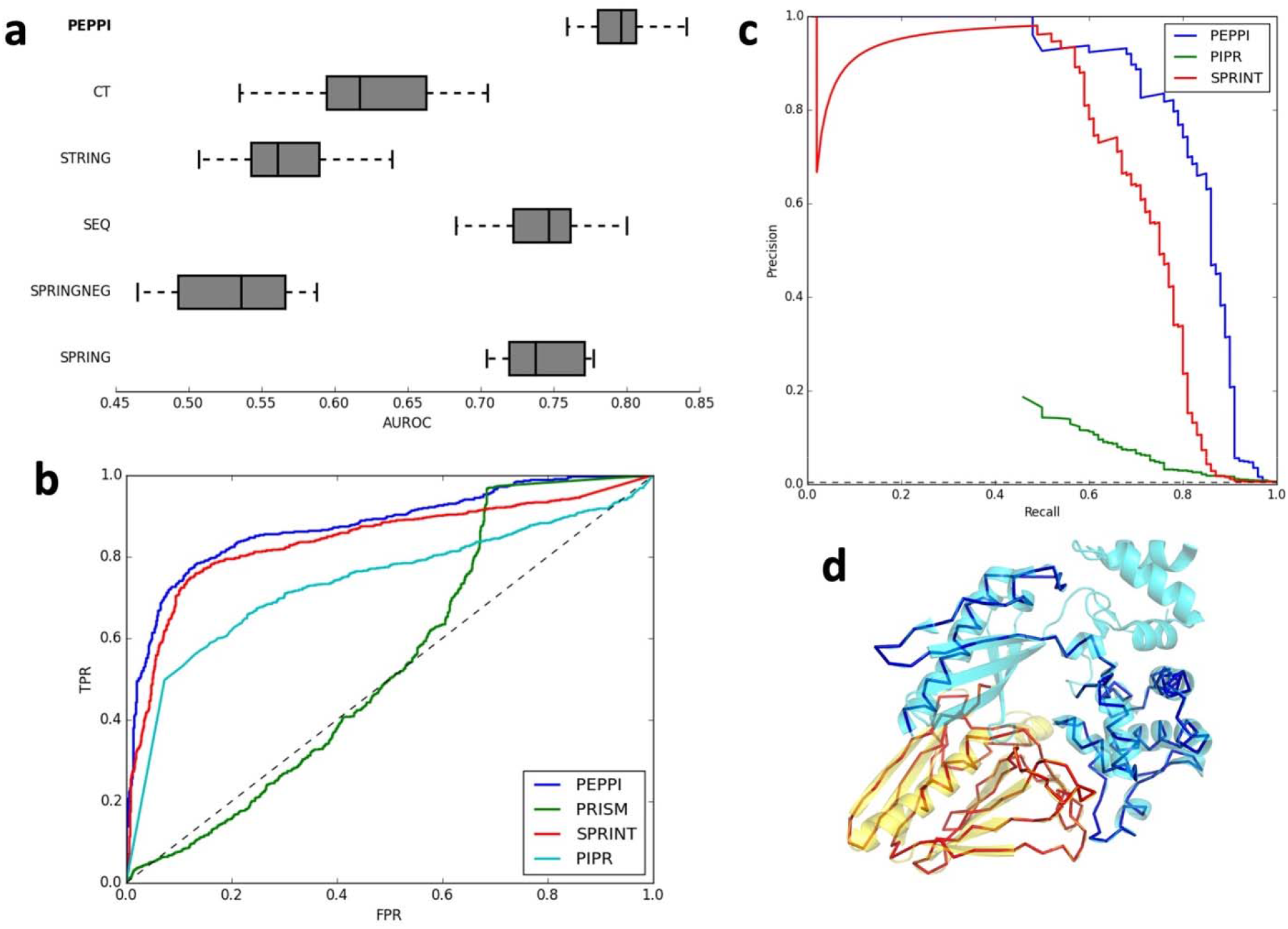
PEPPI benchmark results. (a) 10-fold cross validation AUROC reveals that the full PEPPI pipeline outperforms its component modules: the neural network classifier (CT), the functional association data (STRING), the sequence similarity method (SEQ), the structure similarity method (SPRING), and the non-interaction similarity method (SPRINGNEG). (b) An ROC curve of the performance of PEPPI against PRISM, a structure similarity-based method, SPRINT, a sequence similarity-based method, and PIPR, a deep learning-based method on a balanced testing set. The dotted line represents the performance of random classification. (c) Precision-recall curve of the performance of PEPPI against several other comparable programs on an unbalanced testing set. The dotted line represents the performance of random classification. (d) A superposition of an example dimer model (PDB 3CI0; chain J in red, chain K in blue) on its dimer template structure (PDB 5VTM; chain W in yellow, chain X in cyan).

### 2.2 PEPPI Benchmark and Performance

In order to quantify the classification performance of PEPPI against existing methods, we benchmarked PEPPI alongside PRISM[8], a structure-based similarity predictor, SPRINT[6], a sequence-based similarity predictor, and PIPR[16], a deep learning predictor which utilizes a combination of recurrent and convolutional layers in its architecture. These benchmarks were performed using a randomly selected test set of 798 interacting structure pairs and an equal number of structure pairs involving chains from the same protein complex but known to not come into physical contact (and thus do not form a physical interaction). All structures of this test set were taken from the PDB and were confirmed to have <50% sequence identity to the PEPPI training protein pairs.

The results of the PPI predictions are summarized presented in Figure 2b and Table S1, where it is shown that PEPPI significantly outperforms all competing methods in terms of area under ROC (AUROC), average precision (similar to area under precision-recall curve, AUPRC), and all but SPRINT in maximum achievable Matthew’s correlation coefficient (MCC). PRISM had the lowest performance in this benchmark, which is likely due to its outdated interface structure library, which misses many structures which have been solved since its release to the public. Interestingly, the highest performance from a competing program was seen from SPRINT, a sequence motif-based similarity classifier, and not from PIPR, the more sophisticated deep-learning pipeline. The deep learning architecture of PIPR was optimized based on cross validation performance of species-specific (primarily yeast) interaction datasets; when this method was applied to the species agnostic dataset of this benchmark, PIPR’s ability to accurately classify interactions decreased. As a result, the method which draws its conclusions from explicit similarity (SPRINT) outperforms the method which tries to sub-optimally learn the interaction problem itself (PIPR).

One particular case of interest in this benchmark is the interaction between chain A and chain B of PDB code 1F3M (corresponding to the N- and C-terminal domains of the human kinase PAK1). While this is a true interaction, no hit was found in either the sequence or structure databases after homologous template removal, leading to poor scores for those pipelines (SPRING: 8.937, STRING: not found, SEQ: 0.129, SPRINGNEG: 4.618). However, this case was still classified as positive with a log(LR) of 0.056 due to a high interaction probability from the CT module (0.999), thus demonstrating the utility of CT for rescuing interactions that do not attain significant similarity. On the side of non-interaction classification, the classification of chain G and chain I of 3CJH (corresponding to two Tim13 chains of the yeast Tim8-Tim13 complex) poses an interesting case, as PEPPI was able to classify this as negative (log(LR)=-0.939) where competing programs could not, despite attaining a high SPRING score (34.208), a high CT confidence (1.0), and loose SEQ homology (0.217). The reason for this is a high score from SPRINGNEG (36.108) which pushes down the total interaction likelihood, thus demonstrating a case where SPRINGNEG’s false positive identification ability rescues the pipeline from misclassifying the interaction.

While the previous balanced dataset is convenient for benchmarking, it is not fully reflective of the context in which an interactome prediction algorithm is applied because true interactions are much sparser relative to the total set of pairwise combinations of query proteins in almost all contexts. Therefore, we randomly sampled 100 interacting pairs from the previous test set and paired the 200 chains from these interactions in an all-by-all fashion (excluding homodimer pairings), resulting in an unbalanced test set of 100 true interacting pairs out of 19,990 putative interactions. Due to the high number of interactions, PRISM was excluded from this benchmark because of its slow speed and poor performance on the preceding benchmark. The results of this unbalanced benchmark are presented in Figure 2c and Table S2. The outcome is similar to the previous benchmark, with PEPPI outperforming all other programs, followed most closely by the sequence-based algorithm SPRINT. In this benchmark, however, the superiority of PEPPI is much clearer, as SPRINT is on average more susceptible to false positive detection for comparable recalls. This resulted in a statistically significant difference in max MCC performance between PEPPI and SPRINT, which was not the case on the balanced benchmark set. Also made clearer is the relatively poor performance of PIPR with respect to false positive errors; so many pairs are classified with the highest confidence score that the maximum achievable precision is 0.186. One particular case of interest in this benchmark is the interaction between chain J and chain K of PDB code 3CI0 (part of the type 2 secretion system of enterotoxigenic *E. coli*), an interaction which was detected only by PEPPI. This interaction was able to be detected solely through structural similarity (SPRING: 51.45), with all other modules failing to detect the interaction (STRING: not found, SPRINGNEG: 4.09, SEQ: 0.279, CT: 0.026), leading to a log(LR) of 1.433.

### 2.3 SARS-CoV-2 and Human Interactome Modeling

The COVID-19 pandemic, caused by the SARS-CoV-2 virus, has disrupted the lives of almost every person to some degree, and as of November 2021, over 5 million people have lost their lives to the disease worldwide[23]. SARS-CoV-2 has thus become an essential entity to understand, as our expedient comprehension of this virus translates to the development of therapeutic medicines, such as antiviral drugs and vaccines, for current and future coronavirus infections. One fundamental step towards understanding the function of the virus is to establish what proteins of the host genome each viral protein interacts with. Through modeling the virushost interactome, we can begin to identify the purpose of each viral protein through our functional understanding of the human proteins with which each viral protein interacts. To this end, we have predicted the set of interactions which occur between SARS-CoV-2 and human proteins using PEPPI.

Our SARS-CoV-2/Human interactome model consists of 403 interactions whose likelihood ratios were determined to be greater than 1, i.e., interactions which are more likely to be interacting than not. As shown in Figure 3a-b, the SARS-CoV-2 protein which has the highest number of predicted interactions is the Spike protein (86 interactions), followed by the 2’-O-methyltransferase nsp16 (46 interactions), and the RNA polymerase nsp12 (41 interactions). The highest confidence interaction of this network was the Spike/ACE2 interaction (Figure 3c), which is expected given the extensive study of this interaction due to its essential role in viral entry[24]. PEPPI also correctly predicted Spike to interact with two other host proteins important to viral entry: Furin, a protease which cleaves Spike during entry of SARS-CoV-2 but is not involved in SARS-CoV-1 entry[25], and TMPRSS2, a cell-surface protease involved in viral entry of both SARS-Cov-2 and SARS-CoV-1[24]. The PEPPI results also demonstrated the power of structure similarity-based PPI prediction through the prediction of the PARP15-nsp3 interaction (Figure 3d); this interaction was predicted with high confidence (log(LR)=1.435), mainly due to the SPRING module’s high confidence score (35.6). The PARP proteins are known to interact with the nsp3 macrodomain in other coronaviruses[26], so detection of this interaction in our dataset stands as an important validation.

**Figure 3.**
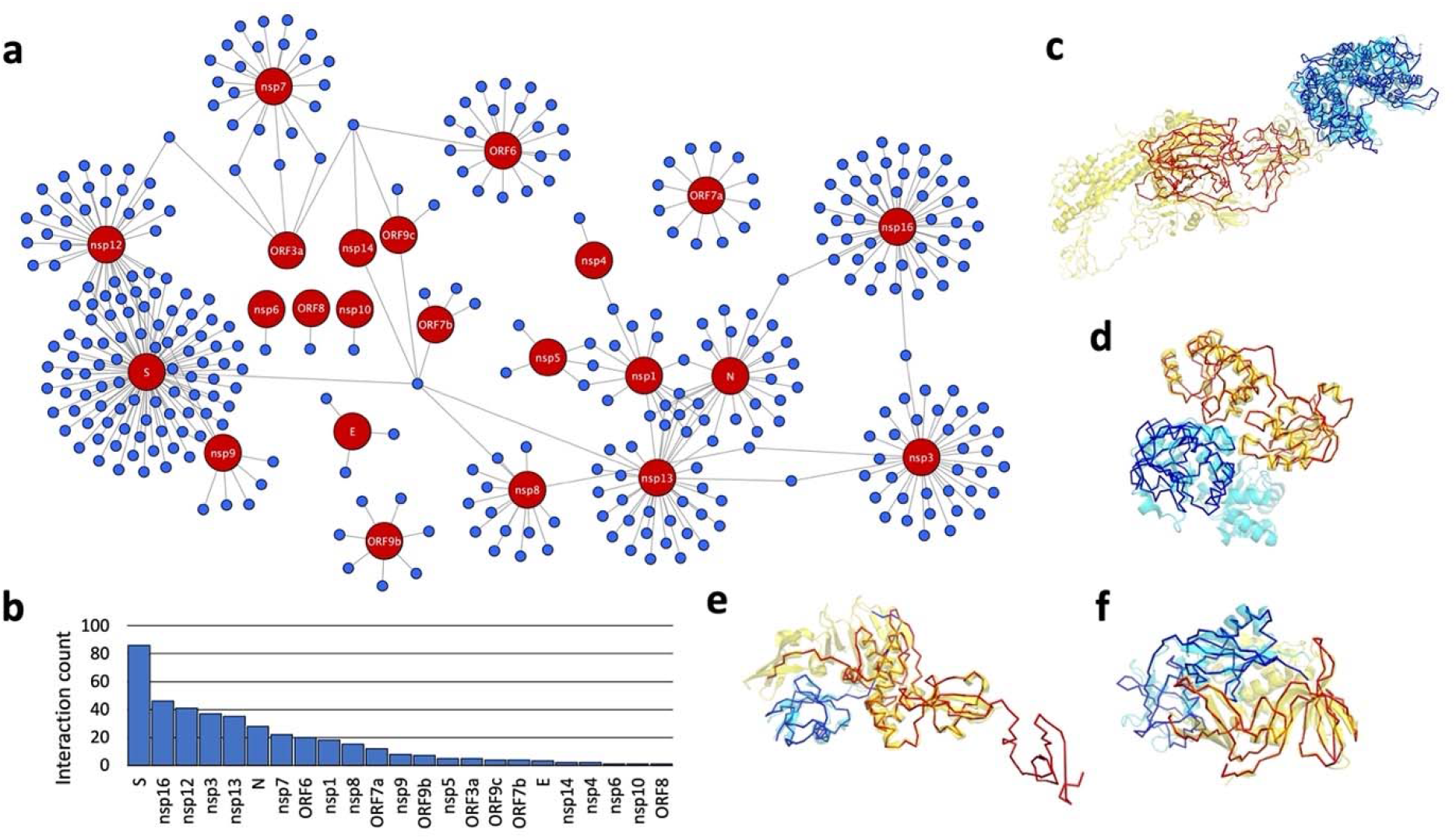
A summary of SARS-CoV-2/human interactome prediction. (a) A network overview of the full interactome of human-covid protein pairs, with SARS-CoV-2 proteins colored red and human proteins colored blue. (b) A bar chart of the number of predicted interactions involving each SARS-CoV-2 protein. Proteins which were not predicted to have any interactions were excluded. (c) A superposition of a dimer model of the top-ranked SARS-CoV-2 Spike (in red) and Human ACE2 (in blue) interaction on its dimer template structure (PDB 6ACG; chain D in cyan, chain C in yellow). (d) A superposition of a dimer model of the SARS-CoV-2 nsp3 macrodomain (in red) and human PARP15 macrodomain (in blue) on its dimer template structure (PDB 2W2G; chain A in cyan, chain B in yellow). (e) A superposition of a dimer model of a domain of the SARS-CoV-2 nsp3 (in red) and human NEDD8 (in blue) on its dimer template structure (PDB 5WFI; chain C in cyan, chain A in yellow). (f) A superposition of a dimer model of a domain of the SARS-CoV-2 nsp3 (in red) and human UBD (in blue) on its dimer template structure (PDB 6BI8; chain C in cyan, chain A in yellow).

To evaluate PEPPI’s overall performance at recapitulating known biology, we constructed a gold standard dataset from PSICQUIC[27] of known interactions for comparison. This dataset consisted of 128 interactions, 94 (73%) of which were predicted by PEPPI. 28 of the 34 missed interactions were not predicted to interact in part because of low SEQ score (<0.3); 19 of these 28 interactions were added to PSICQUIC after the PEPPI databases were constructed, which explains why they were not detected by sequence homology-based approaches. In addition, we compared the overlap between our predicted dataset and a high-throughput experimental dataset recently published in [28]. PEPPI’s predictions only shared one interaction with this dataset (an interaction between MARK3 and ORF9b), but the experimental dataset presents only functional associations due to their use of AP-MS, which is known to pull down entire interacting complexes instead of only the interacting “prey” protein interest. Compared to PEPPI, the dataset in [28] also misses crucial interactions, such as the interactions involving the Spike protein with ACE2, Furin, and TMPRSS2. In fact, only 3 of the 128 (2%) gold standard interactions we isolated from PSICQUIC are present in this dataset. Thus, even in the presence of a high throughput experimental dataset, PEPPI provides a demonstrably useful complement and reveals many direct physical interactions which would otherwise be missed.

Lastly, PEPPI made the potentially significant predictions that nsp3 interacts with the post-translational modifiers NEDD8 and UBD (FAT10). While it is well-documented that the papainlike protease (PLPro) of SARS-CoV-2 nsp3 both deISGylates and deubiquitinates viral proteins to avoid host detection and thus evade immune response[29], there has been less study of the role of the related, ubiquitin-like, post-transcriptional modifiers NEDD8 and UBD in SARS-CoV-2 disease. NEDD8 tags proteins for degradation, has been implicated in the innate immune response to viruses[30], and is a target by some viruses for modulation of host immune response[31]. UBD has been shown to be a ubiquitin-independent and cytokine-inducible modifier targeting proteins for proteasomal degradation[32] and has additionally been shown to have roles in viral infection defense[33]. Furthermore, the top structural templates PEPPI found for nsp3/NEDD8 (Figure 3e) and nsp3/UBD (Figure 3f) were a PLPro in complex with free ubiquitin and a PLPro in complex with the ubiquitin-like and innate-immune-modulating protein ISG15, respectively. We therefore hypothesize that SARS-CoV-2 modulates host innate immune response through interaction of nsp3 with NEDD8 and UBD in a similar manner that nsp3 interacts with ubiquitin.

## 3. Discussion & Conclusion

We have presented a novel PPI prediction pipeline which demonstrates superior performance relative to other approaches. In addition to performance, this method presents a few unique advantages. Firstly, because the structure-based analysis makes use of threading rather than structural alignment, it is much faster than pipelines which need to explicitly model the input chains, while retaining the flexibility of not requiring an input structure. Second, because structure-based analysis is a component of the pipeline, PEPPI can produce rough structural models of the interactions, which can help deepen biological insights such as interface residue determination and can guide follow-up experiments. Finally, because PEPPI is a consensus model, it is not solely dependent on any one methodology to make its predictions; even if all modules classify an interaction with low confidence, if these classifications agree, the final prediction will have reasonable confidence (as we have shown in several examples above). In addition, the consensus classifier is constructed such that if any modules are intentionally excluded or fail to produce a score, a prediction can still be made from the remaining modules.

A few shortcomings and assumptions of the pipeline should also be discussed. First, the interaction predictions made in this pipeline are based largely on similarity to known PPIs, and these modules will only detect interactions with similarity to solved structures or to interactions detected in high-throughput screens. Therefore, the method’s performance will depend on the coverage of our knowledge of the existing interaction space, which is currently far from fully comprehensive. However, this knowledge will expand as more interactions are discovered, so the power of the similarity-based method will improve over time. Second, because PEPPI is similarity dependent, if an interaction is predicted between proteins of two given families, all other proteins in those families will likely also be predicted to interact. In this case, interactions involving proteins of the same families can be sorted by LR; the highest rated interaction is the most likely to be true. Finally, this pipeline predicts the capability for proteins to interact regardless of biological context. As a result, it is possible for some of the interactions predicted here to not exist within the context of the cell due to factors such as incompatible subcellular localization or insufficient expression of the proteins of interest *in vivo*. While this additional biological insight can be useful in pruning the interaction space in a proteome-wide interactome modeling study, it is not explicitly considered in PEPPI’s interaction predictions. Therefore, it is worth validating the interactions predicted with this program with more focused small-scale biochemical studies, such as crosslinking mass spectrometry experiments. Despite these shortcomings, the whole proteome interaction networks modeled by PEPPI can help biologists retrieve existing biology and derive novel biology for their system of interest, as we did for the SARS-CoV-2/Human interaction system. Through the understanding of PPI networks on the whole-proteome scale that PEPPI provides, future studies will be able to better understand the systems-level complexity that underpins biological phenomena as well as target individual edges of the network for therapeutic benefit.

## 4. Methods

### 4.1 Pipeline overview

The PEPPI pipeline performs predictions through a set of independent modules, each of which score the interaction likelihood in their own way. These modules include a conjoint triad trained neural network, a STRING database lookup module, and two “interology” based modules: a threading-based module using a modified version of SPRING and a sequence-based module using BLAST. Scores from each of these modules are transformed into a ratio of likelihood based on pre-trained score probability distributions, and the final likelihood ratio is calculated as the product of likelihood ratios from each independent module (i.e., the sum of log-likelihood ratios). A full description of the pipeline methodology can be found in the Supplementary Material.

### 4.2 SARS-CoV-2 Virus and human host protein sequence collection

The SARS-CoV-2 proteome was collected from the UniProtKB pre-release. Replicase polyprotein 1ab was split into nsp1-16 (excluding nsp11) according to its “chain” regions as described in the “Protein Processing” subsection of its UniProtKB entry; nsp11 was extracted from replicase polyprotein 1a in a similar fashion. As a result, the SARS-CoV-2 sequence set consisted of 31 protein sequences in total. The human proteome, consisting of 20,600 proteins, was also collected from the Uniprot database. All 20600*31=638,600 putative interactions were analyzed with PEPPI; any pairs resulting in a log(LR) greater than 0 were classified as interacting. The “gold standard” dataset is comprised interactions listed in the PSICQUIC database annotated as “direct interaction” as of April 2021 between the proteins of either SARS-CoV-1 or SARS-CoV-2 and human proteins, a total of 128 interactions.

## 5. Acknowledgements

We would like to thank Dr. Gilbert S. Omenn for his consultation and advice in the authorship of this manuscript. Our work was funded by NSF AWD2025426 and NIH AI13467801. The work in this manuscript was performed in part using the Extreme Science and Engineering Discovery Environment (XSEDE), which is supported by the National Science Foundation (ACI1548562).

