## Supplementary Tables and Methods for "PEPPI: Whole-proteome protein-protein interaction prediction through structure and sequence similarity, functional association, and machine learning"

**Table S1.** Performance of the control pipelines against PEPPI in the balanced benchmark. P-values (calculated by 1000-iteration bootstrapping of the distribution of differences in performance between methods) are presented in parentheses.

|  | **AUROC** | **Avg. Precision** | **Max MCC** |
| --- | --- | --- | --- |
| PEPPI | **0.879** | **0.895** | **0.654** |
| PRISM | 0.544 (3.7e-80) | 0.511 (6.2e-115) | 0.373 (6.9e-30) |
| SPRINT | 0.841 (5.3e-7) | 0.853 (6.0e-5) | 0.632 (5.6e-2) |
| PIPR | 0.738 (4.2e-30) | 0.760 (2.7e-24) | 0.469 (5.3e-18) |

**Table S2.** Performance of the control pipelines against PEPPI in the unbalanced benchmark. P-values (calculated by 1000-iteration bootstrapping of the performance difference distribution) are presented in parentheses.

|  | **Avg. Precision** | **Max MCC** |
| --- | --- | --- |
| PEPPI | **0.829** | **0.799** |
| SPRINT | 0.696 (9.3e-4) | 0.729 (1.8e-2) |
| PIPR | 0.122 (4.7e-95) | 0.287 (2.5e-45) |

**Supplementary Methods**

**S1.1 PPI prediction by SPRING threading (“SPRING”/”SPRINGNEG” module)**

SPRING[1] is a program which detects a template dimer structure given a pair of query sequences. However, this program has been altered from its original published state in order to perform better on the task of PPI classification, and therefore, we summarize the new procedure here. First, the pair of query sequences are threaded monomerically by HHsearch[2] through a combination dimer and non-interaction structure library in order to identify monomer structure templates which fit each respective query sequence. Each template structure is scored by the raw alignment score for each query-template alignment; these scores are then Z-normalized by the mean and standard deviation of the top 20,000 templates. A similarity model is constructed for each query chain according to the highest scoring query-template alignment. The top 5,000 template structures (by Z-score) for each of the two chains are paired all-by-all in search of an identical pair of chain IDs in the dimer structure library for SPRING (or the negative structure library for SPRINGNEG), where each identified dimer template pair is scored by the minimum of the two chains’ respective monomeric Z-scores. Once dimer template pairs have been identified, a dimeric model is constructed for the top 100 dimer template pairs by TM-align[3] superposition of the similarity model of each chain onto the dimeric template structure. Each model is then scored according to the SPRINGscore:

$$SPRINGscore=Z+w_{1}TM+w_{2}Dcomplex (1)$$

where *Z* is the Z-score of the dimer template, *TM* is the minimum structure similarity from superposition, *Dcomplex* is the model interface energy as calculated by a modified form of DCOMPLEX[4] which only considers C⍺ atoms, and *w_1_* and *w_2_* are weights with value 4.0 and -0.1 (negative because a lower DCOMPLEX score is more favorable), respectively for SPRING, or 7.6 and 0.0, respectively for SPRINGNEG. The values of these weights were derived on the optimization of 5-fold cross validation performance against a gold standard interaction training set (see “Pipeline training and benchmarking”). The score used in the final classifier is the SPRINGscore of the top-ranked dimer model.

In order to construct the database of template dimer structures, a list of all current Protein Data Bank (PDB)[5] entries was obtained using the database’s RESTful API. For each entry containing at least one protein chain, all “biomolecular assembly” structures were collected. From each biomolecular assembly, pairs of interacting proteins were extracted if both protein chains were at least 30 amino acids long (thus excluding protein-peptide interactions) and if there existed at least 10 interchain contacts (i.e., if there exist at least 10 interchain pairs of C⍺ atoms whose distance is less than 8Å). Each pair is then split into monomeric chains and clustered at 70% sequence identity using CD-HIT[6]. Non-redundant protein pairs are derived from these clusters by pairing each cluster of monomers and extracting one pair at random that exists between the members of each cluster. Additional pairs are extracted if they are structurally non-redundant to the already extracted pairs, i.e. if the new pair has a TM-score < 0.5 by MM-align[7]. Clusters are also paired against themselves to allow for the extraction of homodimeric structures. This procedure resulted in a total of 48,591 non-redundant dimer template structures, constructed from 83,765 monomer chains. In order to make this structure database threadable, an HMM was constructed for each sequence by HHblits[2] using a February 2016 release of the UniProtKB database[8] clustered at 20% sequence identity. The resulting HMM was combined with secondary structure information as determined by DSSP and formatted to be threadable by HHsearch. In addition to the dimer structure library, a negative structure library was constructed from the biomolecular assembly structures, consisting of protein pairs which were found to be in the same complex, but not in physical contact (i.e., no inter-atom distance between C⍺ atoms on differing chains were <8Å apart). These pairs were filtered for non-redundancy in a similar manner to the dimer structure library; however, they were filtered to be non-redundant not only to each other, but also to the original dimer structure library. This way, the negative structure library only consists of pairs which were never shown to interact in any PDB structure. This library consists of 24,568 pairs constructed from 10,708 monomer chains.

In many solved protein complex structures, only the domains which form the interface are present in the structure due to size limitations of the structure solving method. In order to keep this phenomenon from hindering template identification, threading-based domain division of the query is implemented in PEPPI (Figure S1). This procedure collects all query-template alignments above a given Z-score threshold, 8.5, and counts how many of them consist of a large gapped region (>80 AA in length) on one or both termini of the query. If more than half of the total alignments contain a gapped region, the protein is split in two at the optimal domain boundary, and the two domains are analyzed recursively through the same procedure. Recursion stops when either a sequence is found to not have a sufficient proportion of gapped alignments (i.e., is single domain), or to have less than 10 alignments above the Z-score threshold (i.e., is a “hard” threading target). The optimal domain boundary is defined as the average of positions at which the gap region ends (or starts) if only N-terminal (or C-terminal) gaps were found. If both N- and C-terminal gapped alignments are found, each N-terminal boundary is paired with each C-terminal boundary, the midpoint between each pair is determined, and the optimal boundary is defined as the average of these midpoints. For a given pair of query sequences, each domain from one sequence is paired with each domain of the other sequence, and SPRING analysis is done on each domain-domain pair. The highest score derived from domain-domain pairing is kept as the score for the entire query pair.

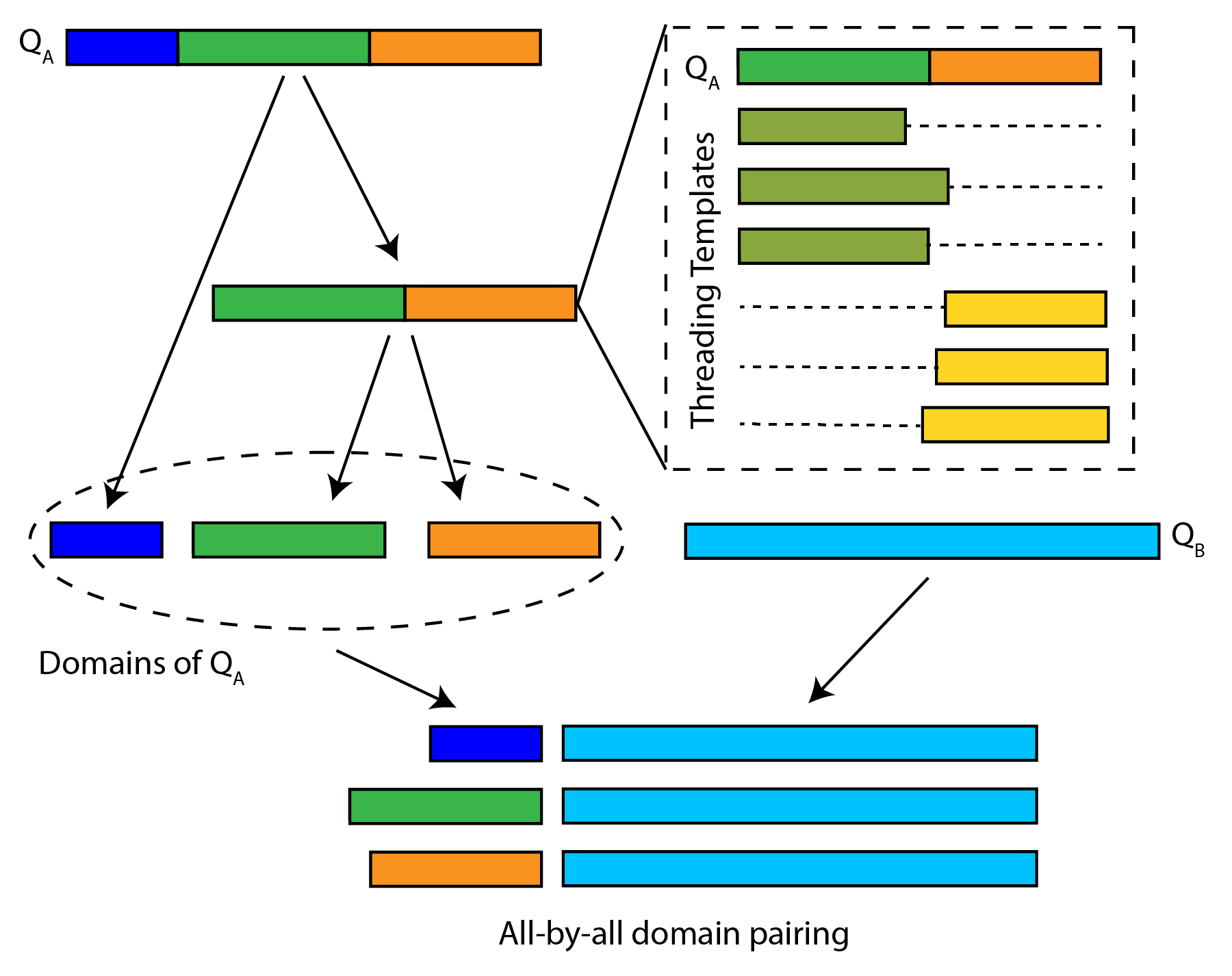

**Figure S1.** An illustration of the domain division procedure for structure-based interology. In this figure, query sequence A (Q­_A_) is divided into domains according to the domain boundary derived from the alignments of its threading templates. After domain division, all domains of sequence A are paired with the single-domain query sequence B (Q_B_) and analyzed individually.

**S1.2 PPI prediction by sequence interology (“SEQ” module)**

The “SEQ” module of PEPPI implements a simple process to identify similar complexes through sequence similarity. The first step of this module is to split apart the two chains of the query and use BLAST[9] to search them through a database of single chain sequences taken from PPIs. The dimer sequence library was constructed from interactions taken from the PSICQUIC service[10]. This service was queried using the “MI:0407 (direct interaction)” term (from the molecular interaction ontology[11]) to retrieve a set of interactions which were annotated to be direct, physical interactions. First, interactions which were not strictly protein-protein interactions were filtered out, including protein-ligand, protein-nucleotide, and protein-complex interactions. Second, amino acid sequences were retrieved for each of the interactions, as they are only presented in PSICQUIC as a set of database identifiers, not as definitive amino acid sequences. Most of this was accomplished using Uniprot’s ID retrieval tool, but sequences were taken directly from the database itself when possible. Finally, the sequences of the dimer structure library were also added to the dimer sequence library. The final dimer sequence library consists of 515,058 interactions in total. It should be noted that this database is redundant, as the same interaction can be represented in multiple databases within PSICQUIC using distinct identifiers. The result of the BLAST search step is a set of single-chain hits sorted by sequence identity to the query. In order to translate this single chain similarity into dimeric similarity, dimers are retrieved from the sets of monomeric hits. The overall score for the dimer is the harmonic mean of the two respective monomeric sequence identities.

**S1.3 PPI Prediction by non-similarity features (“CT”/“STRING”)**

In order to bolster the pipeline’s predictions in the absence of detected sequence or threading hits, PEPPI also uses a neural network model based on the conjoint triad (CT) feature[12]. This feature translates a pair of amino acid sequences into a 686-entry vector based on the frequency of amino acid triads in each of the two sequences. The neural network is implemented as a Multi-Layer Perceptron Classifier in the scikit-learn package[13], with a single hidden layer of size 1000. The model was trained on the dimer structure dataset, as well as an equal number of randomly paired chains. The final probability from this neural network is used in the consensus classifier.

While the primary focus of this pipeline is to predict direct, physical interactions, functional association data can also assist in classification. This functional association data comes from the interaction scores in the STRING v11 database[14]. In particular, the scores used are gene neighborhood (how proximal two genes are in the genome sequence), gene fusion/fission (if other organisms have orthologs of the interacting proteins which have fused into the same polypeptide), co-expression (correlation of gene expression levels across different cell states), and co-occurrence (occurrence of orthologs for each protein in other organisms, regardless of whether they’re known to interact). To retrieve these four scores, the query proteins are monomerically searched through the sequences of the STRING database using BLAST, and any hits with >90% sequence identity are kept. The hits for each chain are then paired against each other and searched through STRING links for a corresponding entry; the four corresponding scores for the first discovered database entry are passed to the classifier. If no hit is found, then missing values are passed to the classifier. It should be noted that the STRING database only contains information for intra-species interactions and does not provide any information on inter-species interactions, such as those in the SARS-CoV-2/Human dataset. PEPPI therefore operates without the STRING module for the SARS-Cov-2/Human application presented in this manuscript.

**S1.4 Pipeline training and benchmarking**

In order to evaluate the interaction likelihood based on the calculated score *x* for a given module in Figure 1, the *x* value is compared against the scores attained by training set consisting of 800 non-redundant interactions from the IntAct[15] database (annotated as “direct interaction” and an MI-score of at least 0.7), and 800 non-redundant non-interactions from the Negatome 2.0[16] database. From the scores of the interaction and non-interaction sets for the given module, two respective probability distributions were created by Gaussian kernel density estimation. From these reference distributions, the likelihood of interaction is expressed as the log-likelihood ratio:

$$log\left( LR \right)=log\left( p\left( x | I \right) \right)-\log\left( p\left( x | NI \right) \right) (2)$$

where *p*(*x|I*) and *p*(*x|NI*) are the conditional probabilities at the score *x* in the interacting and non-interacting reference distributions, respectively. The overall likelihood for a given query pair is the sum of log-likelihoods for all separate modules.

The PEPPI pipeline was then tested on an independent test set consisting of 798 pairs from the dimer structure library, and 798 pairs from the negative structure library, both of which are non-redundant to the pairs in the training set (<50% sequence identity). In order to assess the performance of the pipeline fairly, several non-redundancy measures have been put in place for the benchmark. For the SPRING and SEQ modules of PEPPI and for PRISM, any single-chain templates which have a sequence identity >50% to the query have been removed from consideration. In order to ensure the test set was non-redundant to the training sets of competing programs and that this training set is appropriate for species-agnostic physical interaction prediction, we have constructed a custom training set of 27,967 pairs from the dimer structure library such that they had <50% sequence identity to the test set pairs and an equal number of non-interacting pairs randomly pairing chains from this interaction set. This training set was subsequently used for training SPRINT, PIPR, and the CT modules of PEPPI. Filtering criteria present in PRISM were disabled as the combination of these criteria and our own imposed template non-redundancy rules resulted in a high incidence of no templates being identified. 75 layers were used for PIPR instead of the default 50, as raising the layer number improved 5-fold cross validation performance on the provided training set.
